# The construction and application of a new 17-plex Y-STR system using universal fluorescent PCR

**DOI:** 10.1101/2020.02.18.953919

**Authors:** Jinding Liu, Rongshuai Wang, Jie Shi, Xiaojuan Cheng, Ting Hao, Jiangling Guo, Jiaqi Wang, Zidong Liu, Wenyan Li, Haoliang Fan, Keming Yun, Jiangwei Yan, Gengqian Zhang

**Author notes:** Corresponding author: Dr. Gengqian Zhang, Address: Shanxi Medical University, Jinzhong 030619, Shanxi, China. Additional corresponding author: Dr. Keming Yun, Address: Shanxi Medical University, Jinzhong 030619, Shanxi, China, Dr. Jiangwei Yan, Address: Shanxi Medical University, Jinzhong 030619, Shanxi, China. Jinding Liu and Rongshuai Wang contributed equally to this work.

## Abstract

Y-chromosomal short tandem repeat (Y-STR) polymorphisms are useful in forensic identification, population genetics and human structures. However, the current Y-STR systems are limited in discriminating distant relatives in a family with a low discrimination power. Increasing the capacity of detecting Y chromosomal polymorphisms will drastically narrow down the matching number of genealogy populations or pedigrees. In this study, we developed a system containing 17 Y-STRs that are complementary to the current commercially available Y-STR kits. This system was constructed by multiplex PCR with expected sizes of 126-400 bp labeled by different fluorescence molecules (DYS715, DYS709, DYS716, DYS713 and DYS607 labeled by FAM; DYS718, DYS723, DYS708 and DYS714 labeled by JOE; DYS712, DYS717, DYS721 and DYS605 labeled by TAMRA; and DYS719, DYS726, DYS598 and DYS722 labeled by ROX). The system was extensively tested for sensitivity, male specificity, species specificity, mixture, population genetics and mutation rates following the Scientific Working Group on DNA Analysis Methods (SWGDAM) guidelines. The genetic data were obtained from eight populations with a total of 1260 individuals. Our results showed that all the 17 Y-STRs are human- and male-specific and include only one copy of the Y-chromosome. The 17 Y-STR system detects 143 alleles and has a high discrimination power (0.996031746). Mutation rates were different among the 17 Y-STRs, ranging from 0.30% to 3.03%. In conclusion, our study provides a robust, sensitive and cost-effective genotyping method for human identification, which will be beneficial for narrowing the search scope when applied to genealogy searching with the Y-STR DNA databank.

## Introduction

With the rapid development of DNA analysis technology, STR genotyping methods consisting of multiplex PCR with fluorescently labeled primers and capillary electrophoresis have been conventionally employed in the field of forensic medicine for individual identification and paternity testing(Bai *et al.* 2019; Fan *et al.* 2019).

The Y chromosome acts as a unique tool for forensic investigations since it is inherited through the patrilineal line in a relatively conserved manner(Liu *et al.* 2018a; Tao *et al.* 2019). Unlike autosomal STR markers, for which the typing of two samples must be compared to make a personal identification, typing of Y chromosome markers from stains from a crime scene could be helpful for inferring the potential perpetrator’s origin if his familial DNA typing could be found in a DNA databank, i.e., to find a perpetrator or narrow the investigation scope by the “Sample to family” searching mode(Liu *et al.* 2016; Mo *et al.* 2019; Zhang *et al.* 2019). Y chromosome SNP (Y-SNP) haplogroups are valuable for forensic applications of paternal biogeographical ancestry inference(Lang *et al.* 2019; Ralf *et al.* 2019; Song *et al.* 2019). However, for many of the recently discovered and already phylogenetically mapped Y-SNPs, the population data are scarce in some populations. Y-chromosome microsatellites or short tandem repeats (Y-STRs) were first defined in Europe for forensic purposes and included nine Y-STR loci. Since then, more Y-STR loci have been added to the kits to increase the discrimination power for kinship analysis and human identification and for inferences on population history and evolution(Kayser 2017; Cokic *et al.* 2019). A number of commercial Y-STR kits, such as the Yfiler™ Plus PCR Amplification kit (Applied Biosystems, Foster City, CA, USA), the PowerPlex Y23 System (Promega, Madison, WI, USA), STRtyper-27Y (Health Gene Technologies, China) and the Microreader™ 40Y ID System (Microread Genetics Incorporation, China), are available, and most incorporate 19-40 markers into the single multiplex systems, which have been validated with paternal genetic data in many populations and in forensic casework(Thompson *et al.* 2013; Olofsson *et al.* 2015; Liu *et al.* 2018b). For the purpose of differentiating individuals (only males), Y-STR databases are established for either online public access or within criminal investigation laboratories, which are not available for public access. The US-Y-STR and Y-HRD are established for public access and are used to estimate the Y-STR haplotype frequency or to infer the ethnicity of the donor of a profile(Ge *et al.* 2010; Willuweit and Roewer 2015). The Y-STR databases established in crime laboratories are used to trace male lineages. The famous example of a Y-STR database is the Henan Provincial Y-STR database in China(Liu *et al.* 2016). With this database, the potential profile of a DNA sample from a crime scene is searched by first screening families’ Y-STR haplotypes and then subsequently investigating the identification of the individual in the Y haplotype-matched families. Dozens of cases have been solved in an expeditious manner by using this Y-STR database(Ge *et al.* 2014; Liu *et al.* 2016).

Despite the robustness of the Y-STR markers in these commercial multiplex Y-STR systems and the capacity to discriminate two male individuals in most cases, the coincidence match probabilities are modest compared with a set of standard autosomal STR markers. Henan Province in China has established a large Y-STR (>200000 profiles) database for criminal investigators using either the Applied Biosystems Yfiler or Yfiler Plus PCR Amplification kits(Liu *et al.* 2016). The limited number of Y-STRs has resulted in a few or even large numbers of false-positive matches with the enlargement of the databank. Therefore, improving the discrimination power by increasing the number of Y-STRs would be an effective way to reduce such false matches. Regarding of the strengths and weaknesses of Y-STR markers, more loci are still required to provide more information to assist forensic investigations, as also emphasized by Ge and Liu et al(Ge *et al.* 2014).

In this study, we developed a new 17 Y-STR typing kit, which exceeds the current Y-STR system containing the trinucleotide loci DYS718 and DYS719; the tetranucleotide loci DYS715, DYS709, DYS713, DYS607, DYS708, DYS723, DYS712, DYS605, DYS726 and DYS722; and the pentanucleotide loci DYS716, DYS714, DYS717, DYS721 and DYS598(Zhang *et al.* 2004a; Zhang *et al.* 2004b; Zhang *et al.* 2012).

Currently, the analysis procedure is often performed by evaluating fluorescent fragments via capillary electrophoresis (CE)(Wang *et al.* 2019; Xu *et al.* 2019). Universal fluorescent PCR is a cost-effective genotyping method in which fluorescent fragments are obtained with non-labeled forward primers, non-labeled reverse primers and fluorescent universal primers by a two-step amplification(Blacket *et al.* 2012; Oka *et al.* 2014; Asari *et al.* 2016).

The combination of Y-STR and an economic fluorescence labeling technique can reduce the expenditures for forensic laboratories. We then used a new kind of genotyping method featuring FAM/JOE/TAMRA/ROX-labeled universal primers (M13 (−21) universal primer 5’-TGTAAAACGACGGCCAGT-3’)(Oka *et al.* 2014). We validated this kit according to the recommendations on forensic analysis proposed by the International Society of Forensic Genetics(Gill *et al.* 2001; Gusmao *et al.* 2006). Following these guidelines, we defined the Y-STR and allele nomenclature and assessed the kit for PCR conditions, sensitivity, accuracy, species specificity and the effects of DNA mixtures and mutation rates. Our results showed that the 17-plex Y STR typing system is a reproducible, accurate, sensitive and economical tool for forensic identification.

## Materials and Methods

### DNA samples

The following procedures were performed with the approval of the Ethics Committee of Shanxi Medical University. Written informed consent was obtained from all participants.

A total of 1600 blood samples were collected from 930 unrelated male samples, 10 unrelated female samples and 330 unrelated father-son pairs. The 1590 male samples were collected from 724 males (330 unrelated father-son pairs, which were confirmed by using autosomal STR analysis, and 64 unrelated males) in Taiyuan City (Shanxi Province), 162 unrelated males in Chongqing City, 154 unrelated males in Ulanqab City (Mongolia), 155 unrelated males in Sanmenxia City (Henan Province), 95 unrelated males in Foshan City (Guangdong Province), 113 unrelated males in Hainan Li, 63 unrelated males in Hainan Miao and 124 unrelated males in Jingzhou City (Hubei Province) (see Figure S1). The 10 female samples were recruited from the Shanxi Han population.

DNA was extracted by using a QIAamp^R^ DNA Investigator Kit (QIAGEN China (Shanghai)) and was quantified by using a Quantifiler^R^ Human DNA Quantification Kit (Applied Biosystem, USA) on a Bio-Rad Real-time PCR System (Bio-Rad Laboratories, USA). The quantification process was performed per the manufacturer’s instructions.

The control DNAs 9948, 2800M, and 9947A were purchased from Promega (Promega, Madison, WI, USA).

The species specificity study evaluated the capacity for the system to avoid the detection of genetic information from nontargeted species. Nonhuman samples from chickens, cattle, fishes, pigs, rabbits, rats, sheep and shrimps were obtained from Shanxi Medical University Animal Center. DNA was extracted using a QIAGEN DNA Tissue Mini Kit (Qiagen, USA) and was quantified by the standard OD260 method. For each species, 10 ng of DNA was amplified by the 17-plex Y-STR assay following the standard protocol. The human DNA sample 2800M (4 ng) was used as a positive control.

### Primer design

The loci were selected from our previously published articles(Zhang *et al.* 2004a; Zhang *et al.* 2004b; Zhang *et al.* 2012). Sequences for each locus were obtained from NCBI (https://www.ncbi.nlm.nih.gov/) using a standard nucleotide BLAST (Basic Local Alignment Search Tool) search. We selected 17 autosomal Y-STRs, which were classified into four sets (Sets A, B, C and D) based on the needs of multiplex amplification.

An 18-base universal M13 (−21) sequence (TGTAAAACGACGGCCAGT) was used as the standard sequence. The universal tail was added to the 5’ termini of the forward primers in each set. The primers were labeled with four fluorescent dyes (FAM, JOE, TAMRA and ROX) according to the four sets. All primers were synthesized by Shanghai Sangon Biological Engineering Technology & Services Company, Shanghai, China.

### PCR amplification

All individuals were analyzed in four independent multiplex PCRs (Sets A, B, C, and D). Each PCR was performed by using 1 ng of DNA in a total volume of 15 μL, containing (0.008 μM-0.8 μM) forward primers with universal tails, reverse primers, 0.25 μM of the fluorescent universal M13(−21) primer and 1x PCR MasterMix (Mei5 Biotechnology (Beijing) Co, Ltd). The reaction conditions were as follows: 95°c for 10 min; 25 cycles of 95°c for 25 s, 56°c for 25 s, and 72°c for 25 s; 8 cycles of 95°c for 25 s, 53°c for 25 s, and 72°c for 25 s; and a final extension at 72°c for 60 min. PCR products (0.4 μL of each set) were added to 10 μL of HiDi™-Formamide (Applied Biosystems), containing 0.3 μL of the GeneScan™HD 500 ROX™ Size Standard (Applied Biosystems), and were detected by capillary electrophoresis (CE) using an ABI PRISM^R^ 3130 Genetic Analyzer (Applied Biosystems) with a 36-cm capillary and performance-optimized polymer (POP4). Data were analyzed using GeneMapper ID software (Applied Biosystems). Signal intensities were represented by relative fluorescence units (RFU). The bins and panels for the multiplex system were programmed for genotyping.

Allelic ladders were created for the 17 loci by using TA cloning to represent the range of alleles observed in the Shanxi population(Zhou and Gomez-Sanchez 2000; Yao *et al.* 2016). Alleles were amplified independently at each locus with monoclonal cells, and the products of each locus were then diluted and reanalyzed to produce a single allelic ladder for each locus. Finally, those single allelic ladders were mixed in appropriate proportions to create a “cocktail”. Each reaction (total volume=100 μL) contained 0.2 μM of each locus-specific primers, 2 μL of monoclonal cells and the reagent from the multiplex PCR kit (Mei5 Biotechnology (Beijing) Co, Ltd) at 1x concentration. The reaction conditions were as follows: 95°c for 10 min; 25 cycles of 95°c 25 s, 58°c 25 s, and 72°c 25 s; 8 cycles of 95°c 25 s, 53°c 25 s, and 72°c 25 s; and a final extension at 72°c for 60 min.

### Sensitivity

The sensitivity study estimated the capacity for the system to obtain the whole profile from a range of DNA quantities. It was performed with control DNA 9948 (Ori-Gene company, USA), and total DNA inputs were prepared in a serial dilution to 1 μL with the following template amounts: 12 ng, 4 ng, and 0.4 ng (3 ng, 1 ng and 0.1 ng DNA in each reaction). All experiments were carried out according to the parameters provided above.

### DNA mixtures

Male and female DNAs (formed by 2800M and an anonymous female sample) were mixed in four combinations, which were previously diluted to 1:1, 1:50, 1:100, and 1:500. For each combination, the male DNA was maintained at 1 ng, and the amount of the female DNA was varied from 1 ng to 500 ng in each reaction. For male and male mixtures, a total of 2 ng of DNA was input in each reaction. The samples were prepared using an anonymous male sample and 2800M human genomic DNA, with mixture ratios of 1:0, 1:1, 1:2, 1:4 and 1:9. Each mixture was tested in triplicate to reduce the accidental error and to ensure the accuracy of the results analysis.

### Statistical analyses

The allele and haplotype frequencies were counted directly. Gene diversity (GD) was calculated as: 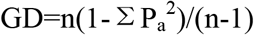, where n is the total number of samples and P_a_ is the relative frequency of the a^th^ allele at the locus, respectively(Nei 1973). Haplotype diversity (HD) were calculated as 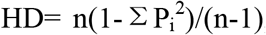, where P_i_ is the frequency of the i^th^ haplotype and n indicates the total number of samples(Siegert *et al.* 2015; Li *et al.* 2020). The discrimination capacity (DC) ratio was calculated as 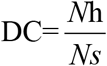, where Nh represents the number of unique haplotypes and Ns represents *Ns* the total number of individual samples(Li *et al.* 2020). Population Fst genetic distance and P value (P value was corrected by Sidak’s correction for multiple testing (P<0.0018, 28 pairs)) were obtained using Arlequin software to assess the genetic structure(Zhu *et al.* 2014). PCA based on the genotypes of the 17 Y-STRs in the eight populations was performed with SPSS 22.0. The subpopulation structure was examined via model-based clustering algorithms implemented in STRUCTURE 2.3.4, which was based on a Bayesian Markov Chain Monte Carlo algorithm. Analyses were performed from K=2 to K=8 using the no-admixture model and correlated allele frequencies to estimate the 17 Y-STRs. Structure Harvester was applied to estimate the optimal K value.

The mutations were counted directly, and the locus-specific mutation rate was calculated as the number of observed mutations divided by the number of father-son pairs. The 95% confidence intervals (CIs) were estimated from the binomial probability distribution available at http://statpages.info/confint.html(Wu *et al.* 2018).

### Quality control

The above experiments were performed in the Forensic Genetics Laboratory of the School of Forensic Medicine of Shanxi Medical University, P.R. China, which is an accredited laboratory, in according with quality control measures. The DNAs 9947A and 9948 were used as negative and positive controls, respectively. All the alleles observed in this assay were validated by Sanger sequencing.

#### Data Availability Statement

The reagent, software, and data are available upon request. The authors affirm that all data necessary for confirming the conclusions of the article are present within the article, figures, and tables. Besides, we have uploaded supplementary material to figshare.

## Results

### Assessment of the system

The 17 loci were divided into four panels, and each panel was assigned a color dye. The allelic ladders and internal size standard in the 17-plex system are shown in Fig. 1. The set of designed primers act according to unified parameters, with a consistent temperature, to achieve a stable and balanced amplified sensitivity and efficient amplification in multiplex PCR processes. Primer-related information on each locus (sequence, melting temperature and distance from repeat motif, concentration, PCR product size and dye label) is listed in Table 1. In all, we successfully established the 17-plex Y-STR system with universal fluorescence-labeled primers. The genotype results of 9948 and 2800M are shown in Table 1.

**Fig.1.**
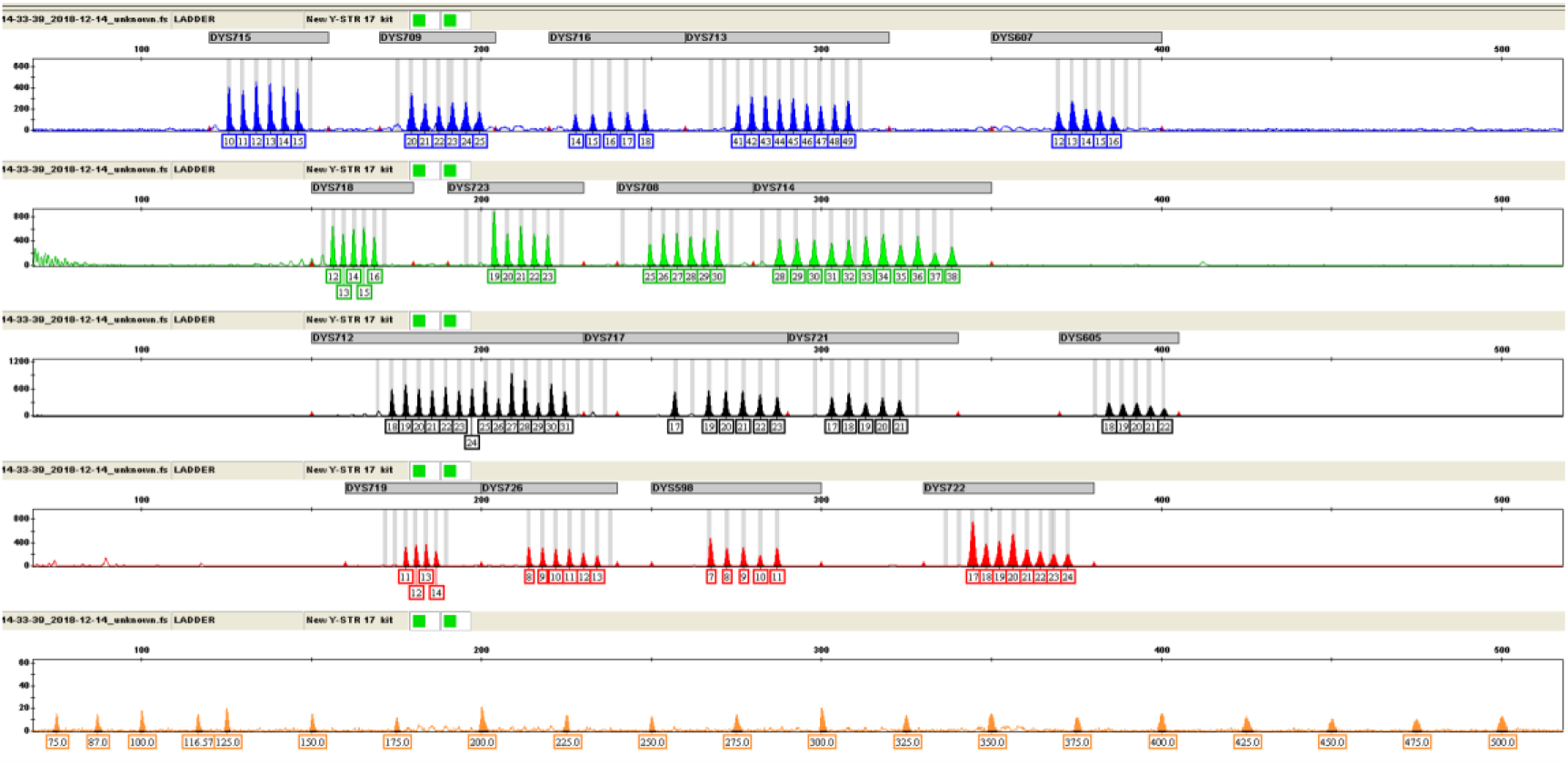
Electropherogram of allelic ladders and internal size standard in the 17-plex system. The four dye panels for the allelic ladders correspond to (from top to bottom) FAM (blue), JOE (green), TAMRA (yellow), ROX (red) dye-labeled peaks. The genotype is shown with the allele number displayed underneath each peak. The fifth panel shows the internal size standards labeled with Orange500 dye (a total of twenty fragments:75, 87, 100, 116.5, 125, 130, 175, 200, 225, 250, 275, 300, 325, 350, 375, 400, 425, 450, 475 and 500bp).

**Table 1.**
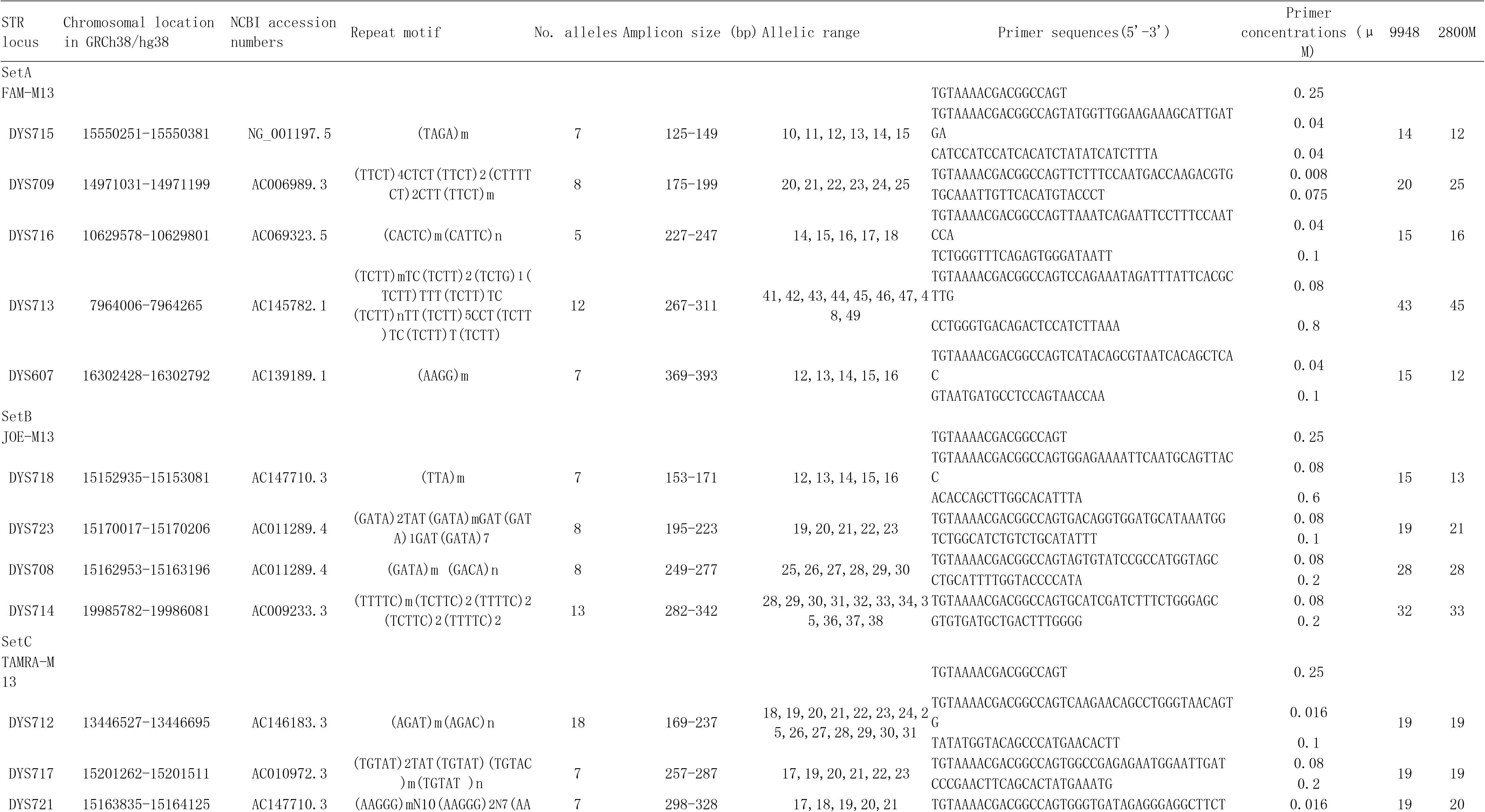
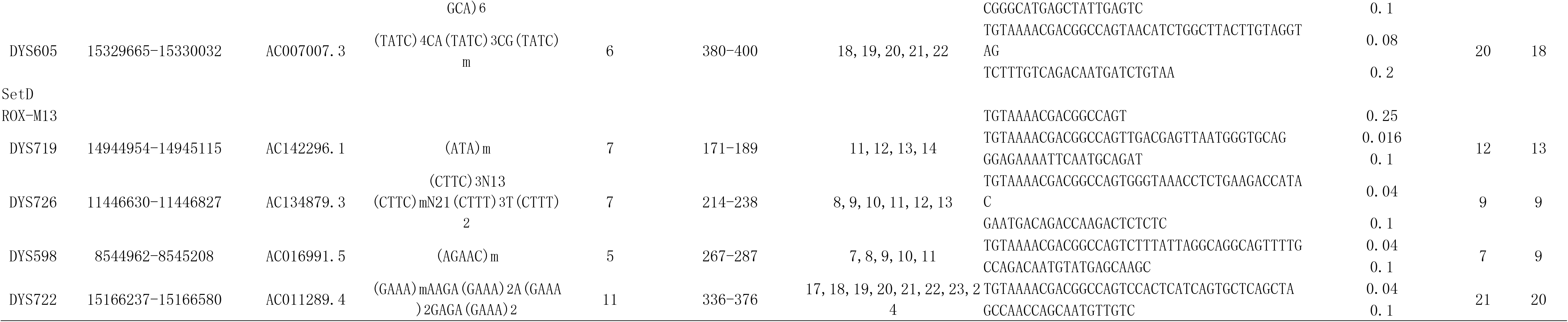
General information on loci used in the new 17 Y-STR system

Mixed samples consisting of DNA from two or more individuals are frequently encountered in many forensic cases. Thus, we assessed the typing system’s capacity to analyze DNA mixtures. Mixtures of two individuals (male 2800M and an anonymous female sample) were examined in various ratios (1:0, 1:1, 1:10 and 1:100) with 1 ng of template DNA in each set for male/female mixtures. Testing was performed in triplicate to ensure the accuracy of the results. All peak heights of the male 2800M could be called for ratios of 1:1 and 1:100 (Fig. 2a). The two male samples (an anonymous male sample and 2800M) were used to evaluate male/male mixtures with ratios of 1:0, 0:1, 1:1, 1:2, 1:4 and 1:9 for a total of 2 ng of template DNA in each set. The minor alleles dropped out at the 1:4 DNA (the anonymous male sample) ratio. The peak height ratios in the 1:4 and 1:9 mixtures were also 10% or less. We were unable to call alleles for some of the minor profiles at the DYS716 locus in the 1:4 ratio samples and at the DYS718, DYS712 and DYS605 loci in the 1:9 ratio samples (Fig. 2b). These results indicate that the 17 Y-STR system can be used to genotype DNA samples in mixtures with relatively small mixture ratios.

**Fig.2.**
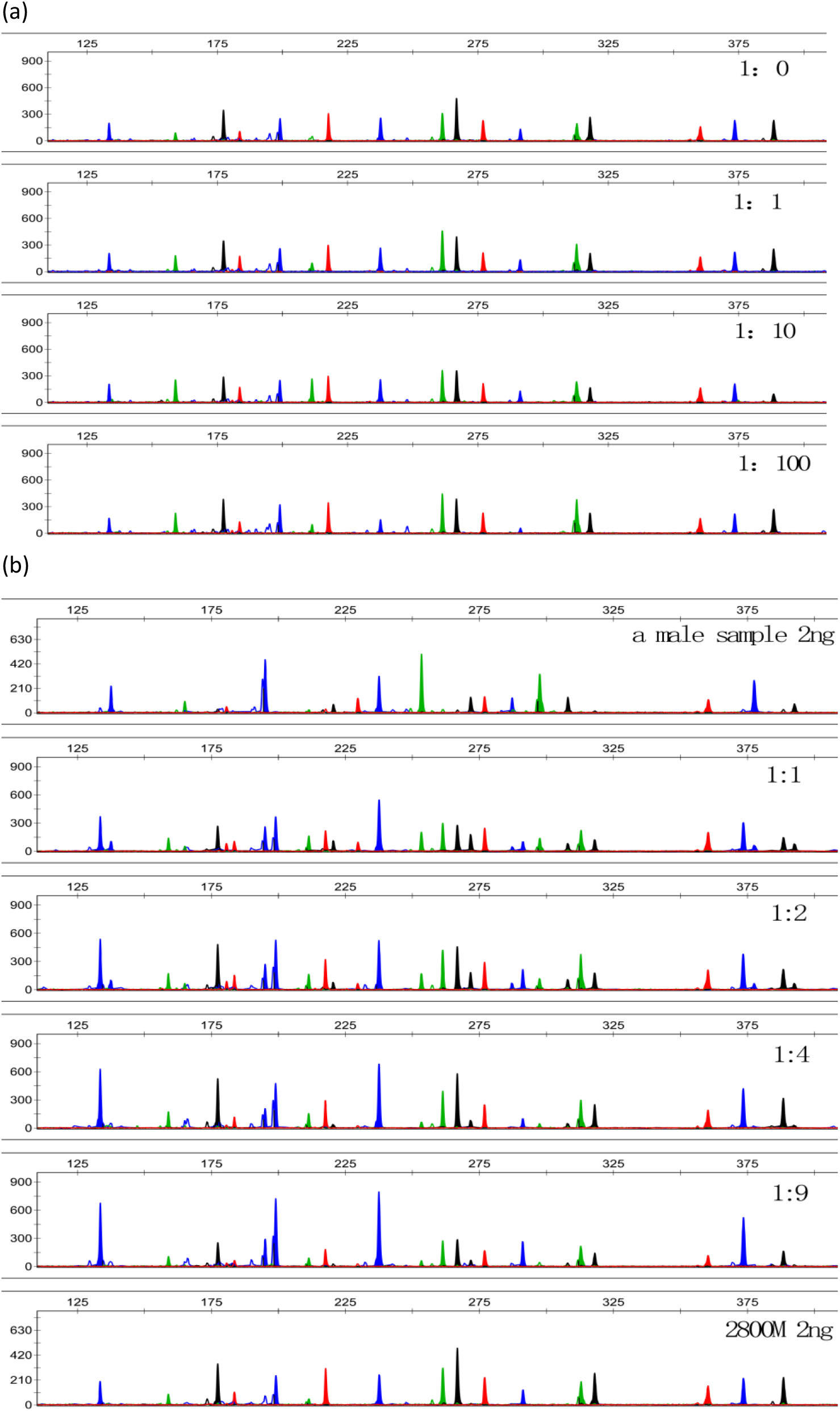
Representative electropherogram of DNA mixture. (a) was formed by 2800M (1ng DNA template in each set) and an anonymous female sample at the ratios 1:0, 1:1, 1:10, 1:100. (b) Mixtures of two male samples (an anonymous male sample and 2800M) were examined in various ratios (1:0, 0:1, 1:1, 1:2, 1:4 and 1:9) with 2 ng total template DNA in each set.

We assessed the sensitivity of the reaction by setting 50 RFU as the limit of detection and 60% as the stochastic threshold for peak balance using varying amounts of 9948 control DNA. We were able to obtain the full profile of sample 9948 when the DNA template provided was greater than 0.04 ng. For samples with more than 0.04 ng, the peak height increased with increasing amounts of DNA template in the reaction. Allelic drop-out and allelic imbalance occurred when the amount of template DNA was reduced to less than 0.04 ng. We thus used 4 ng of template DNA for our typing system (Figure S2).

Nonhuman genomic DNA samples from common animal species (chicken, cow, fish, pig, rabbit, rat, sheep and shrimp) and female samples were amplified using the 17-plex Y-STR system. The results did not reveal the presence of any allele peaks within the genotyping range (Figure S3).

On this basis, it was concluded that the developed STR system was robust and unlikely to be affected by the presence of genetic material from these animal species and female samples.

### Population data

We used the 17-plex Y-STR system to genotype 1260 unrelated Chinese males recruited from eight Chinese ethnic groups (Table S1). Table S2.1 shows the allele frequencies in eight populations. The allele frequencies ranged from 0.0025 to 0.9919. A total of 143 alleles were detected using this system. Table S2.2 shows the allele frequencies and parameters in 1260 individuals in China. A total of 1255 distinct haplotypes were generated; 1250 individuals had unique haplotype profiles, while the remaining 10 individuals exhibited the same haplotype as one other individual. Gene diversity (GD) ranged from 0.2939 (DYS726) to 0.9176 (DYS712). The haplotype diversity (HD) and discrimination capacity (DC) were 0.999993696 and 0.996031746, respectively (see Table S3).

STRUCTURE software (the program STRUCTURE implements a model-based clustering method for inferring population structure using genotyping data consisting of unlinked markers) runs for K=2-8 showed an optimum K value (K=5) estimated by Structure Harvester (http://taylor0.biology.ucla.edu/structureHarvester/). A bar plot of K=5 analysis reveals that the Hainan Miao and Hainan Li samples can be primarily separated from other Chinese Han populations (Fig. 3a). The PCA demonstrated the population stratification in the tested samples (Fig. 3b).

**Fig.3.**
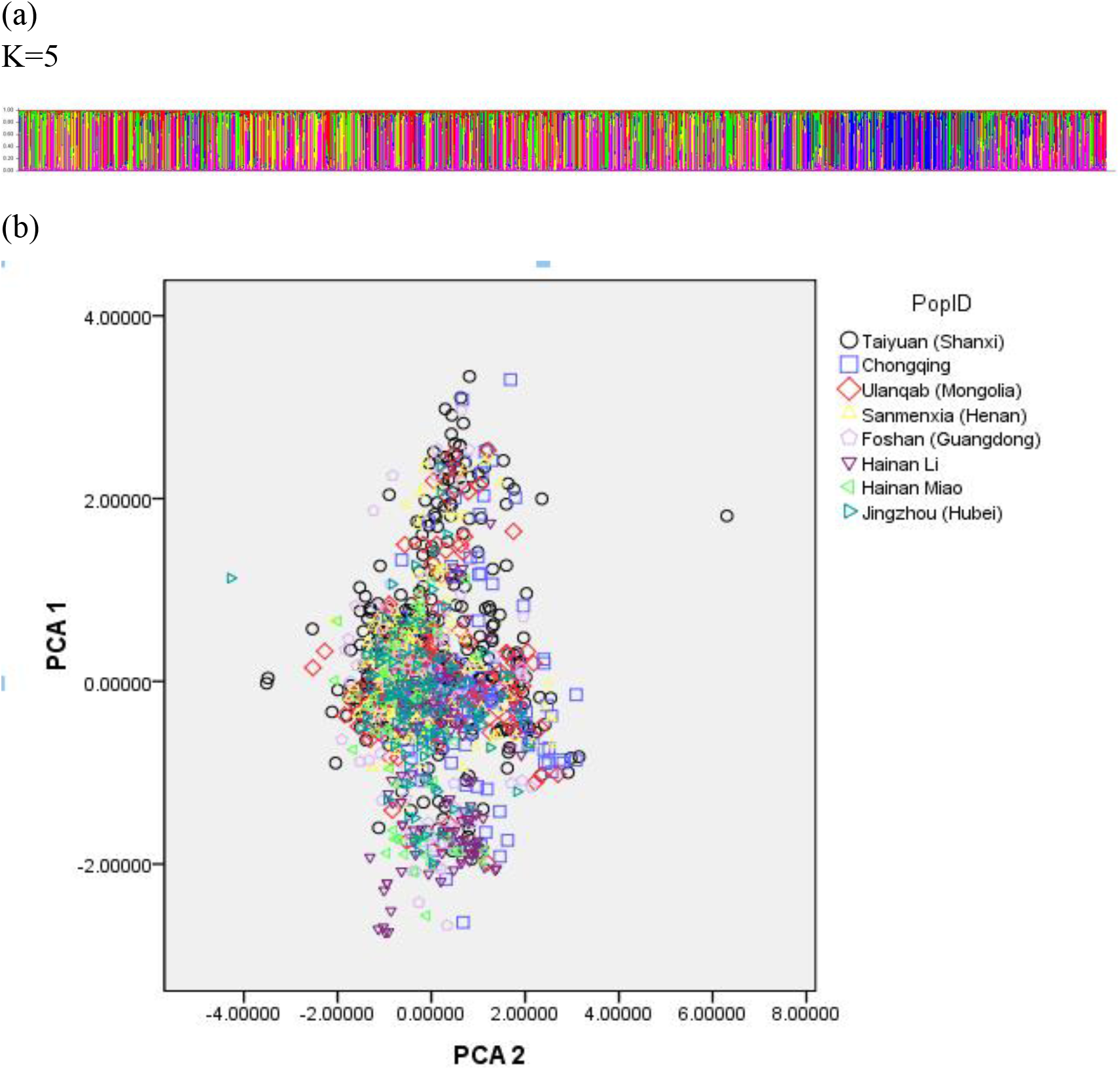
Component analysis results for 1260 Chinese samples from 8 population with the 17 selected Y-STR markers. (a) Bar plot of STRUCTURE run for K=5. The bars with different colors represent different areas of samples. (b) 2-dimensional plot of PCA analysis. The first two principal components (PC1 vs. PC2) demonstrate the population stratification in the tested samples.

The Fst and P values are indicators for evaluating the distribution differences of allele frequencies among populations. The Fst and P values for pairwise interpopulation comparisons were calculated based on allele frequencies of 17 Y-STRs by analysis of molecular variance (AMOVA) performed with ARLEQUIN version 3.5 software between eight populations (see Table S4). There were significant differences after Sidak’s correction for multiple testing (P<0.0018, 28 pairs) at DYS716, DYS713, DYS607, DYS718, DYS714, DYS717, DYS721, DYS605, DYS719, DYS726 and DYS598. At DYS716 and DYS713, the Chongqing population was different from the other seven populations. At the DYS607 locus, the Taiyuan (Shanxi) population could be differentiated. At DYS718, the Hainan Li population could be differentiated from the other seven populations. At the DYS714, DYS721, DYS605, DYS719, DYS726 and DYS598 loci, the Jingzhou (Hubei) population could be differentiated. At the DYS717 locus, the Hainan Li and Jingzhou (Hubei) populations could be differentiated from the other six populations.

A total of 330 allele transmission events from 330 father-son pairs were analyzed at the 17 Y-STRs. Mutations were observed in 38 father-son pairs in DYS715, DYS709, DYS716, DYS713, DYS607, DYS718, DYS714, DYS723, DYS712, DYS721, DYS719, DYS726 and DYS598 (see Table 2). The average mutation rate was 0.89%. The Y-STR DYS712 had the highest mutation rate of 3.03%. Highly polymorphic STRs had high mutation rates. However, no mutations were found in DYS708, DYS605, DYS717 and DYS722, which had relatively high gene diversity values. There were 38 mutations observed, including 16 repeat gains versus 22 repeat losses (1:1.375) and 35 one-step mutations versus three two-step mutations (11.667:1). A one-step mutation is shown in Figure S4.

**Table 2.** Mutation rates of 17 Y-STRs in a Han population of Shanxi, China

| Y-STR locus | Allele<br>transmission | Number of<br>mutation | Number of<br>gains | Number of<br>losses | Number of two steps<br>mutation | Mutation<br>rate | Binominal 95%<br>CIs |
| --- | --- | --- | --- | --- | --- | --- | --- |
| DYS715 | 330 | 1 | 1 |  |  | 0.30% | 0.0001–0.0168 |
| DYS709 | 330 | 1 |  | 1 |  | 0.30% | 0.0001–0.0168 |
| DYS716 | 330 | 4 | 2 | 2 | 1 | 1.21% | 0.0033–0.0307 |
| DYS713 | 330 | 2 | 1 | 1 |  | 0.61% | 0.0007–0.0217 |
| DYS607 | 330 | 4 | 1 | 3 |  | 1.21% | 0.0033–0.0307 |
| DYS718 | 330 | 2 |  | 2 |  | 0.61% | 0.0007–0.0217 |
| DYS714 | 330 | 5 | 3 | 2 | 1 | 1.52% | 0.0049–0.0350 |
| DYS723 | 330 | 1 |  | 1 |  | 0.30% | 0.0001–0.0168 |
| DYS712 | 330 | 10 | 4 | 6 |  | 3.03% | 0.0146–0.0550 |
| DYS721 | 330 | 1 | 1 |  |  | 0.30% | 0.0001–0.0168 |
| DYS719 | 330 | 3 | 1 | 2 | 1 | 0.91% | 0.0019–0.0263 |
| DYS726 | 330 | 1 |  | 1 |  | 0.30% | 0.0001–0.0168 |
| DYS598 | 330 | 3 | 2 | 1 |  | 0.91% | 0.0019–0.0263 |
| Total |  | 38 | 16 | 22 |  | 0.89% |  |

## 4. Discussion

In this study, we developed a cost-effective multiplex 17 Y-STR genotyping method using four fluorescent universal primers. Multiplex PCRs using ≧ 4 ng of DNA produced intense signals for all 17 loci, all genotypes from eight different populations were fully concordant with STR profiles, and the mutation rate was assessed with 330 father-son pairs.

Standard DNA profiling using sets of well-selected, largely standardized, highly polymorphic autosomal STRs is very suitable for identifying the donor of a single-source crime scene trace, as long as the person’s STR profile already exists in the DNA database. Currently, the STR profiles obtained from crime scene traces are compared with the stored profiles in the forensic DNA database to look for a match. However, this comparative autosomal STR profile matching for human identification is not successful for completely unknown perpetrators, whose STR profiles are not yet available. In addition, if the traces are from multiple-source stains, the autosomal STR profiles can be compromised because of the masking effect of the main contributor. As in sexual assault cases, for example, based on the excess of epithelial cells from the female major contributor, it is often difficult to single out the autosomal STR profile of the male contributor. This is where Y-chromosome STR profiling comes into play, as only the male perpetrator, but not the female victim, carries a Y chromosome.

However, when dealing with specific cases, additional loci are required. For cases to discriminate related males belonging to the same paternal lineage or to separate paternal lineages in populations with low Y-chromosome diversity, loci with relatively high mutation rates would be useful to improve the power of discrimination. For paternal lineage discrimination, loci with relatively low mutation rates would perform better. Therefore, more Y-STR loci should be studied.

In this study, we used a cost-effective multiplex genotyping method with four fluorescent universal primers. There are two different designs for labeling the PCR amplicons. One approach involves the addition of universal tails to only forward (or reverse) primers, while the other adds the tails to both forward and reverse primers. To reduce interactions between primers, we used the first strategy in this study. We selected four fluorescent labels with different colors (FAM, JOE, TAMRA and ROX), which are commonly used by the Health STRtyper series. This choice also enabled us to use the same capillary electrophoresis running module (Dye Set Any5dye) with the 36-cm capillary and POP4 polymer.

The usefulness of multiplex PCR systems has often been discussed based on interloci balances. By optimizing the concentrations of the fluorescent universal primers, our system can produce well-balanced signals of the 17 Y-STR loci. The concentrations of the reverse primers are generally 10 times higher than those of the forward primers when increasing the production of fluorescent fragments, which is higher than Masaru Asari et al recommended in their research(Asari *et al.* 2016). Optimal PCR conditions are required to acquire a robust detection system. To obtain well-balanced signals among the 17 loci, we evaluated the concentration ratios of the fluorescence-labeled primer, forward primers and reverse primers. We found that differential primer concentrations yielded much better results than those using even primer concentrations. The best signals were obtained when the ratios of the forward/reverse primer concentration ranged from 1:1 to 1:10. The PCR was performed with higher annealing temperatures for 25 cycles to amplify the target genome DNA and lower annealing temperatures for eight cycles to integrate the fluorescence to the PCR products. For commercial STR kits, the cost of fluorescence labeling cannot be neglected. Using the optimized PCR parameters, we were able to decrease the cost of fluorescence labeling to 25% for the following CE detection.

Previous studies of the 17 Y-STRs identified 90 alleles(Zhang *et al.* 2004a; Zhang *et al.* 2004b; Zhang *et al.* 2012). In this study, we detected 143 alleles in 8 populations. For all 17 Y-STRs, only the allele frequencies of DYS607 and DYS598 had been investigated in Cape Muslim, Belgium and southeast China populations(Shi *et al.* 2009; Cloete *et al.* 2010; Claerhout *et al.* 2018). The number of alleles was the same in our population. We evaluated the mutation rates in all 17 loci in this study, while only DYS607 was evaluated in the Belgium population. The STRUCTURE result estimated the classification relationships among populations. A bar plot of K=5 revealed that our 8 populations could not be separated into 8 clusters, which meant that our system could be used in all Chinese populations. PCA is a classical nonparametric linear dimensionality reduction technique that extracts the fundamental structure of a dataset without the need for any modeling. It can be used for inferring population clusters and assigning individuals to subpopulations. It is not possible to significantly distinguish all of the samples. Fst and P values were used to demonstrate the genetic differentiation. After Sikad’s correction, the P values at 11 loci were less than 0.0018, suggesting that these markers had large differences in gene frequencies between the two populations.

STRs with mutation rates greater than 10^−2^ were termed rapidly mutating Y-STRs (RM Y-STRs)(Ay *et al.* 2018). In our study, 38 mutations were observed in 13 loci: 35 mutations were one-step mutations, and 3 mutations were two-step mutations. Nine loci (DYS715, DYS709, DYS713, DYS718, DYS723, DYS721, DYS719, DYS726, DYS598) showed low mutation rates (<10^−2^), and four loci (DYS716, DYS607, DYS714 and DYS712) could be regarded as RM Y-STRs, with the highest mutation rate of 3.03 ×10^−2^ at DYS712 and the lowest mutation rate of 1.21 ×10^−2^ at DYS716 and DYS607. The mutation rate at DYS607 in our population (1.21 ×10^−2^) was higher than that in the Belgium population (2.54 ×10^−3^)(Claerhout *et al.* 2018). Among the 17 Y-STR markers, four loci (DYS708, DYS717, DYS605 and DYS722) with no mutations were observed, nine loci (DYS715, DYS709, DYS713, DYS718, DYS723, DYS721, DYS719, DYS726 and DYS598) had mutation rates lower than 1.00 ×10^−2^, and the rest of the loci (DYS716, DYS607, DYS714 and DYS712) had mutation rates higher than 1.00 ×10^−2^ in our population. The 13 highly polymorphic Y-STR markers could be used to improve the discrimination capacity in populations with low genetic diversity. The newly discovered 4 Y-STR loci with high mutation rates can be used to differentiate different individuals from the same paternal lineage.

This multiplex assay remains a supplemental DNA tool which could provide additional Y-STR information. With the increment of 17-plex Y-STRs application in the future, we proposed to amplify in a single reaction and labeled the one of the two primers for each Y-STR at the 5’ end with dye to decrease the amount of input DNA.

## Conclusion

With recent developments in forensic science, Y-STR analysis applied for forensic purposes has been continually improved. This article developed a new 17-plex Y-STR typing system for forensic genetic testing that incorporated the loci from of the current Yfiler Plus kit. Developmental validation studies that included PCR conditions as well as testing the cross-reactivity, sensitivity, anti-interference, and stability of the method and population data analysis have demonstrated it to be a sensitive, robust, and highly informative tool for use in forensic casework.

## Acknowledgments

This work was supported by Chongxin Judicial Expertise Center (Hubei) & Liu Liang Personal Studio (CXLL20170002) and Program for the Top Young and Middle-aged Innovative Talents of Higher Learning Institutions of Shanxi.

